# Submodular sketches of single-cell RNA-seq measurements

**DOI:** 10.1101/2020.05.01.066738

**Authors:** Wei Yang, Jacob Schreiber, Jeffrey Bilmes, William Stafford Noble

**Affiliations:** Department of Genome Sciences, University of Washington; Department of Electrical and Computer Engineering, University of Washington; Paul G. Allen School of Computer Science and Engineering, University of Washington

## Abstract

Analyzing and sharing massive single-cell RNA-seq data sets can be facilitated by creating a “sketch” of the data—a selected subset of cells that accurately represent the full data set. Using an existing benchmark, we demonstrate the utility of submodular optimization in efficiently creating high quality sketches of scRNA-seq data.

---

By capturing variation of gene expression within a population of cells, single-cell RNA-seq (scRNA-seq) measurements add an additional dimension to already large gene expression datasets. As a result, scRNA-seq datasets can be massive. For example, a recent scRNA-seq analysis of 61 staged mouse embryos yielded measurements of *>*2 million cells. ^1^ Clearly, meta-analyses that aim to aggregate scRNA-seq data from multiple such studies will be challenging.

One common strategy to facilitate analysis of very large datasets is to identify and remove redundant examples. In scRNA-seq, this strategy can be used to select a subset of the cells in an experiment that show different patterns of gene expression. The selected subset, sometimes referred to as a “sketch” of the full dataset, can then be analyzed using clustering or cell type assignment methods. ^3^ A recently described method for selecting sketches of scRNA-seq data, Geosketch, is based on minimizing a particular distance function—the Hausdorff distance—between the full dataset (the “ground set”) and the sketch. ^8^

Here, we propose *submodular optimization* as a theoretically-grounded and powerful framework for selecting a sketch of scRNA-seq data. Loosely speaking, submodular optimization can be considered a discrete analog of convex optimization, where the goal is to identify a set of discrete elements, rather than a collection of continuous values, that optimize an objective function.

Submodular functions are set functions that satisfy the property of diminishing returns: if we think of a function *f* (*A*) as measuring the value of a sketch *A* that is a subset of a larger set of data items *A* ⊆ *V*, then the submodular property means that the incremental “value” of adding a data item *s* to a sketch *A* decreases as the size of *A* grows (e.g., *f* (*A* + *s*) − *f* (*A*) ≥ *f* (*B* + *s*) − *f* (*B*) whenever *A* ⊆ *B* and *s* ∉ *B*). Unfortunately, searching for a sketch of maximal quality, as measured by *f* (*A*), is computationally infeasible for an arbitrary set function; however, when the set function is submodular, then the quality can be approximately maximized (i.e., the quality of the identified solution is a least constant factor times optimal) in low-order polynomial time ^6;17;19^. Moreover, the approximation ratio achieved by these optimization algorithms is provably the best achievable in polynomial time, assuming *P* ≠ *NP*. For these reasons, submodular optimization has a long history in economics, ^2;27^ game theory, ^24;25^ combinatorial optimization, ^5;16;23^ electrical networks, ^18^ and operations research. ^4^ Furthermore, submodular optimization has recently been used with great success for selecting sketches of text documents, ^12–14^ recorded speech, ^15;29;30^ image compendia, ^26^ sets of protein sequences, ^11^ sets of genomics assays, ^28^ and sets of genomic loci. ^7^

The Hausdorff distance that Geosketch minimizes is not submodular but can be viewed as a robust form of a commonly used submodular function (Supplementary Note 1). Hence, we hypothesized that switching to a submodular optimization framework, with its fast algorithms and performance guarantees, would yield better (or at least as good) sketches much more quickly than Geosketch.

To test this hypothesis, we applied two submodular optimization toolkits—an open source Python package called “apricot” ^22^ and a commercial tool provided by Summary Analytics Inc. (smr.ai)—to the four benchmark datasets that were analyzed in the Geosketch paper. To evaluate the methods, we used the same performance measure employed by Hie *et al*., namely, the count of the number of cells of the rarest cell type that are included in the sketch. Here, the intuition is that a good sketch is one that undersamples common cells that presumably occupy dense regions of the transcriptional space. We observe that, in all four datasets, the sketches produced by the submodular approaches outperform Geosketch, often by a large margin (Figure 1A). Both submodular approaches use roughly the same objective function, though the commercial tool includes proprietary forms of the user-specified objective. These modifications explain the overall better performance of smr.ai relative to apricot on these evaluation tasks. The submodular approaches also compare favorably to Geosketch in running time (Figure 1B). The apricot software, which is implemented in Python using both numpy ^20^ and numba ^10^ to accelerate computation, performs comparably to Geosketch for smaller selected sets but runs more slowly as the size of the selected set gets larger. On the other hand, the smr.ai tool, which is implemented in C++, is generally an order of magnitude faster than Geosketch.

**Figure 1:**
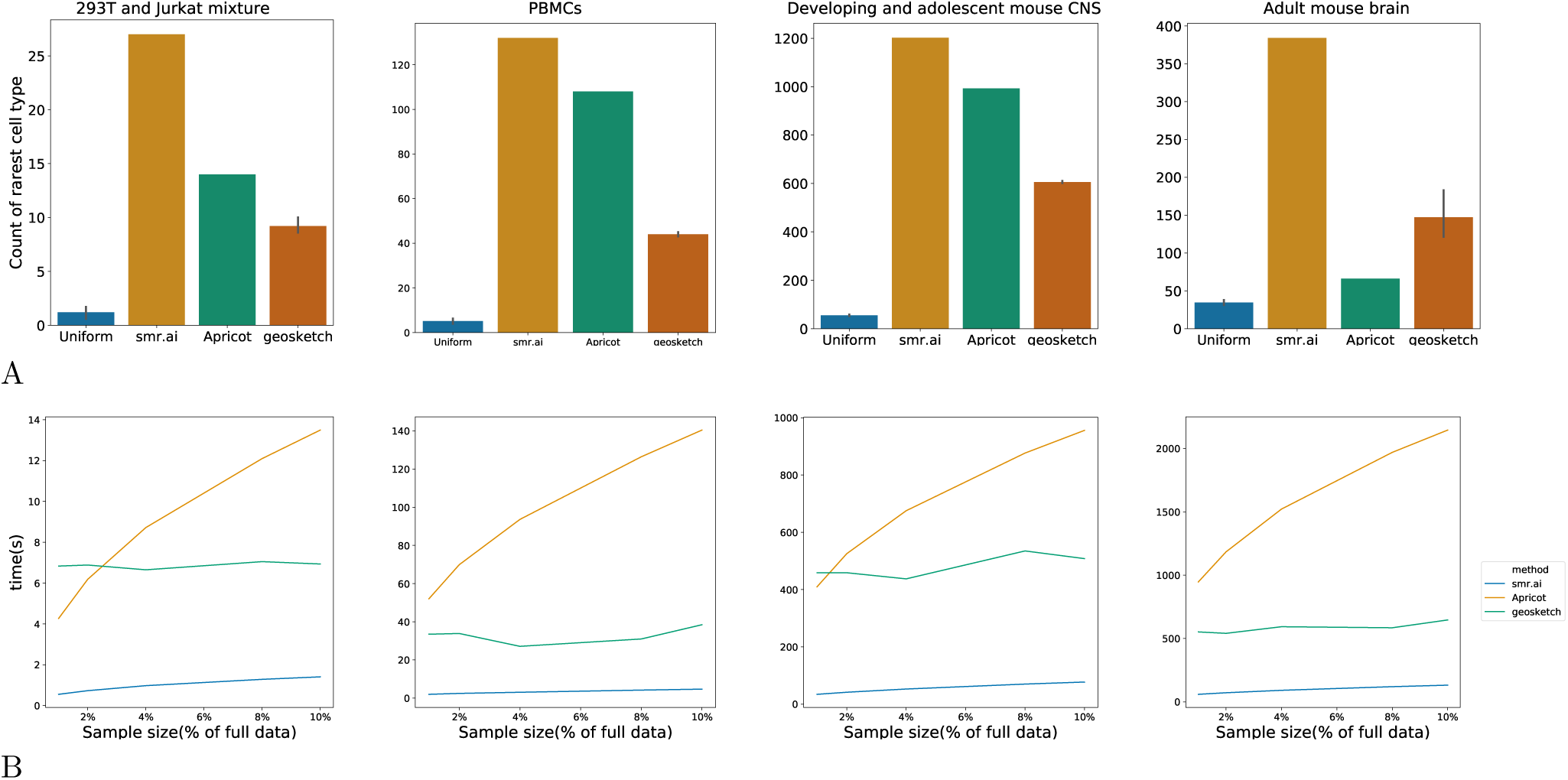
(A) Each panel plots, for a specified benchmark dataset, the number of cells of the rarest type that were selected by each of the four sketching methods. (B) Each panel compares the wall clock times required by Geosketch, apricot and smr.ai to produce sketches of varying sizes.

Overall, our empirical results suggest that submodular optimization provides a powerful and efficient way to summarize large-scale scRNA-seq datasets. Of particular note, the results we report here have not been optimized with respect to parameter selection—they represent the very first objective function that we tried. The space of submodular functions is large and very diverse, and particular functions can in principle be constructed to obtain particular types of sketches. Thus, users of these tools may wish to experiment with varying the form of the submodular objective to obtain sketches that are, for example, particularly enriched in rare cell types or that pay particular attention to capturing outliers.

## Methods

### Data

We downloaded four sc-RNA-seq datasets previously used to assess the performance of Geosketch: a 293T and Jurkat mixture dataset with 4,185 cells, ^32^ a peripheral blood mononuclear cell (PBMC) dataset with 68,579 cells, ^32^ a developing and adolescent central nervous system (CNS) dataset with 465,281 cells, ^31^ and an adult mouse brain dataset with 665,858 cells. ^21^ For each dataset, we focused on the same rarest cell type as Geosketch: 28 293T cells (0.66% of the total number of cells in the dataset) in the 293T and Jurkat mixture dataset, 262 dendritic cells (0.38%) in the PBMC dataset, 2,777 ependymal cells (0.60%) in the mouse CNS dataset, and 1695 macrophages (0.25%) in the adult mouse brain dataset.

### Submodular optimization

We ran both apricot and smr.ai using a “feature-based” objective ^9^ of the form

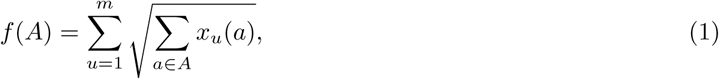

where *X* is a dataset represented as an *n* × *m* matrix with columns *x*_*u*_ for feature *u, u* is the index of a single feature in the dataset, *m* is the number of features in the dataset, and *x*_*u*_(*a*) is the value of feature *u* for example *a*. The square root function is critical, since its concavity is what provides a diminishing returns property. Note that, in general, this type of feature-based selection requires non-negative feature values (i.e., *x*_*u*_(*a*) ≥ 0). In the case of scRNA-seq data, the read counts are naturally non-negative. For both smr.ai and apricot we subsequently normalized the read counts for each cell to a unit vector, maintaining the non-negativity of each feature. Prior to analysis by apricot, the expression values of each gene were linearly rescaled to the range [0, 1]. Feature-based selection was carried out by calling apricot’s `FeatureBasedSelection` function with default parameters.

The apricot software is freely available under MIT license at https://github.com/jmschrei/apricot. The smr.ai software can be, on request, made to be used freely by academic researchers via http://smr.ai.

### Geosketch

Geosketch was imported as a Python package obtained from http://cb.csail.mit.edu/cb/geosketch. Each data matrix was reduced to a dimensionality of 100 PCs by randomized PCA before the sketch, as implemented in Geosketch. The matrix of 100 PCs and subset size were then passed to the `gs gap` function to generate a list of indices of selected cells. Note that we consider the randomized PCA part of the Geosketch method. Hence, in contrast to the results reported in Hie *et al*., the error bars in Figure 1A reflect variation from both the randomized PCA and the Geosketch algorithm, rather than just the Geosketch algorithm.

### Timing

To compare running times, we generated sketches of varying sizes on a single thread on a 2.70GHz Intel Xeon CPU E5-2680 with 250 GB of memory. Wall clock times for apricot and Geosketch were recorded using the Python “time” module. For smr.ai, we used the wall clock time reported by the software. As is common in the high-performance computing literature, we repeated each timing test 10 times and reported the minimum wall clock time, to reduce any possible effect of operating system interference.

## Author Contributions

Data analysis was carried out by WY, JS, and WSN. The manuscript was written by WY and WSN, and was edited by all authors

## A Supplementary Note 1

The goal of Hie *et al*. ^8^ is to minimize the Hausdorff distance between the fixed ground set (they call it 𝒳, we call it *V*) and the subset (they call it *S*, but we use *A* ⊆*V* above). Hence, *V* is fixed and only *S* ⊆*V* is variable, and their goal is to perform the following optimization:

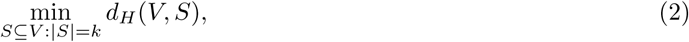

where *d*_*H*_ (*T, S*) = max_*t*∈*T*_ min_*s*∈*S*_ *d*(*t, s*) for arbitrary subsets *T, S* ⊆ *V*. This expression is parameterized by element-pair distances, i.e., *d*(*t, s*) for *t, s* ∈ *V*. This optimization in Equation (2) is equivalent to the problem

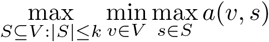

where *a*(*v, s*) = *α* − *d*(*v, s*) is an affinity (or similarity) between pair *v, s* and where *α* is a positive constant. If we define

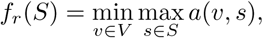

then this objective is a minimization over a set of submodular functions, since the function *g*_*v*_(*S*) = max_*s*∈*S*_ *a*(*v, s*) is submodular in *S* for all *v* ∈ *V*. A well-known standard submodular function known as the facility location function has the form

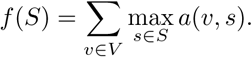

When we compare *f*_*r*_(*S*) with *f* (*S*) we see that *f* (*S*) is a sum over the set of submodular functions *{f*_*v*_(*S*)*}* while *f*_*r*_(*S*) is the minimization over the same set of submodular functions. Hence, Geosketch, which maximizes *f*_*r*_, maximizes the worst case over the set *{f*_*v*_(*S*) *}* of submodular functions. In contrast, when using the facility location function, we maximize the average case over the same set of functions. Unfortunately, the minimization over a set of submodular functions does not preserve submodularity so we cannot use the same fast algorithms while being afforded the same mathematical guarantee. Hence, at least until the space of submodular functions for the problem of finding sketches of scRNA-seq measurements has been fully investigated, it seems that attempting to maximize the worst case may be premature. Our results above, showing good results using a simple feature-based objective, support this suggestion.

